# A simple and viable approach to estimate population trends

**DOI:** 10.1101/2022.03.18.484446

**Authors:** Mario Schlemmer

**Affiliations:** University of Klagenfurt

**Keywords:** ecological stability, temporal variability, correlation, trend strenght

## Abstract

Changes in population abundances over time are of central concern for environmental conservation and the understanding of population dynamics. The standard slope estimator has restricting assumptions and lacks a constant range, which makes its interpretation less intuitive. Herein, more robust measures of trend. It is based on proportional difference and can be used for data on various scale types. If it is applied to ranks it yields the correlation coefficient. Related measures of association are described that assess the relationship between species, including a rank-order correlation that is sensitive to top ranks and a correlation for continuous data that is more robust than the correlation coefficient. All proposed measures have a range between –1 and +1. Furthermore, they can provide a common ground for evaluating trend strength and strength of association for populations undergoing very different dynamics.

## Introduction

Measures of trend strength that evaluate changes in population abundances over time are an important tool in ecologists’ toolkit. A standard measure of trend strength is based on the slope of the regression line for a linear regression(Gerrodette, 1987; White, 2019). The measure has several limitations. The metric has no constant range, which makes the values hard to interpret. Different formulas also yield differently standardized values, making results difficult to compare. It is also based on the assumption of linearity, population growth or decline has to be constant and trends associated with nonlinear population dynamics are not measured appropriately. Here I introduce a new measure of trend strength based on proportional differences that address these limitations before I describe related measures of association.

## Measuring trend in population abundances

Abundance can refer to the number of individuals at a site, biomass, or coverage. A new measure of trend strength can be calculated from proportional differences between groups of values and the mean abundance. Beginning with time step 2, the abundance at each time step is added to the abundances at all later time steps and divided by the mean abundance 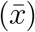, then the number of observations included in each comparison(c) is subtracted. The proportional difference for each of the *n* – 1 comparisons contributes a positive or negative value to the sum of differentials D that is standardized by division through the maximum difference MD, which is calculated the same way as D, but for values that are ordered from lowest to highest.

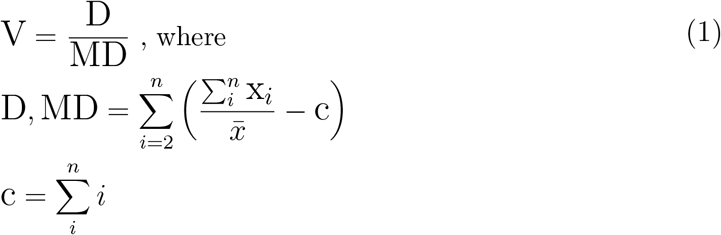

Bounded between −1 and 1, V measures monotonic population trends. Given that there is at least one difference between abundances, the value will be 1 if abundances stay the same or increase consistently over time, and −1 if they stay the same or decrease consistently. If no monotonic association exists between abundances and time steps the value is 0.

**Figure 1:**
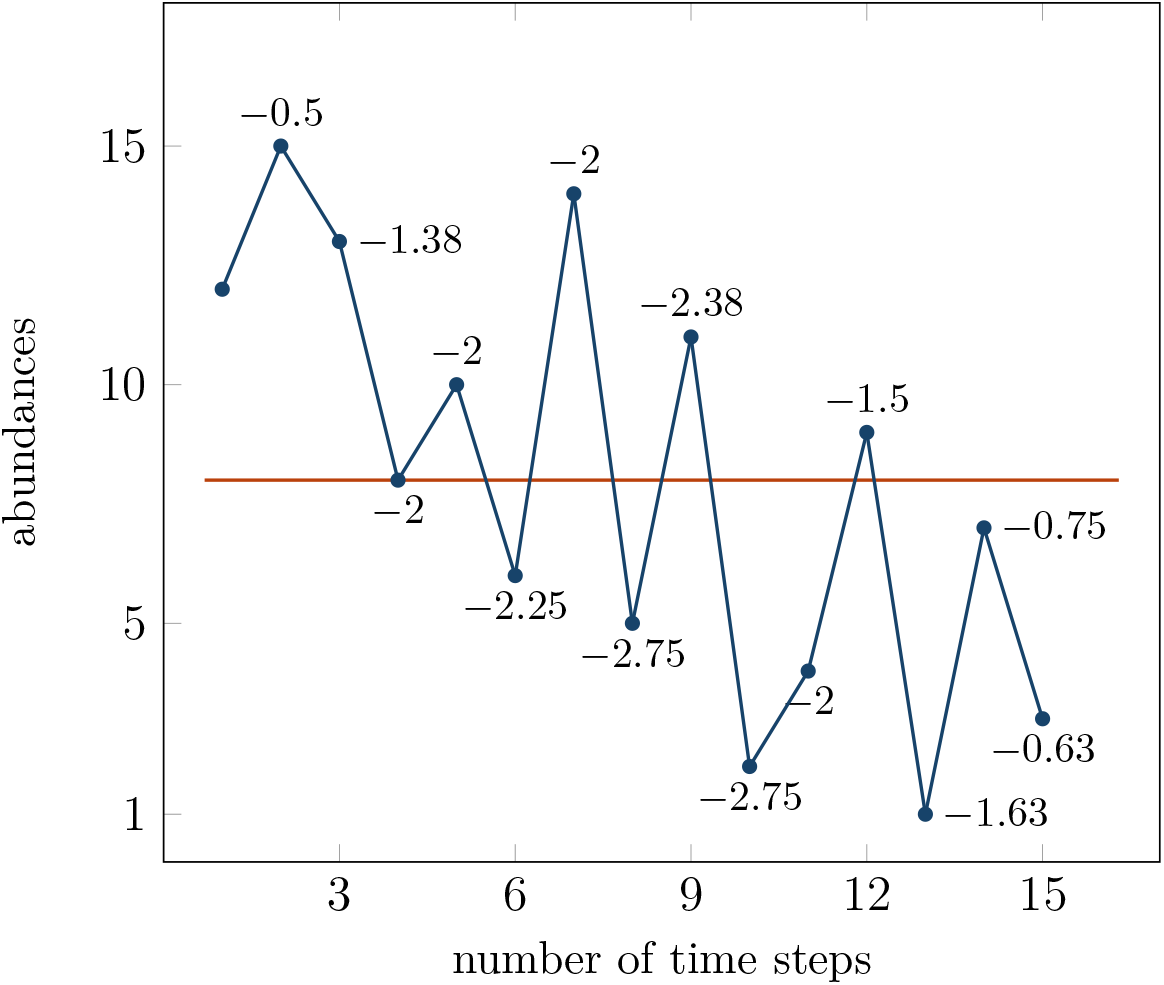
Proportional difference for sets of abundances at each time step(blue) relative to the mean abundance(red).

Monotonic relationships can accurately measure linear trends and various non-linear trends that are equally relevant to ecology. In the context of population abundances, this includes exponential population growth if ideal environmental conditions allow fast population growth, but also logistic growth associated with populations that reach the carrying capacity of the environment. Through simulations, I found that V has a high degree of concordant validity with the slope coefficient, but it places a stronger weight on marked differences between the first and last time step. The strongest negative trends as measured by V have higher abundances at the first time step and low abundances at the last time step for a negative trend, compared to the slope coefficient. Similarly, high positive values have lower abundances at the first time step and higher abundances at the last. A certain degree of sensitivity to initial and final abundances is an appealing property in a trend measure.

V can be used for the evaluation of trends for data of different scale types: nonnegative metric data, rank data, ordinal data, and binary presence/absence data. If used for environmental variables that take on negative values the mean may be zero. In that case, a small constant like 1 × 10^−9^ can be added to the mean to avoid indetermination. A worked example that illustrates the calculation of V is presented in Table 1, where *x_i_* refers to the population abundances for a fictional dataset.

**Table 1.**
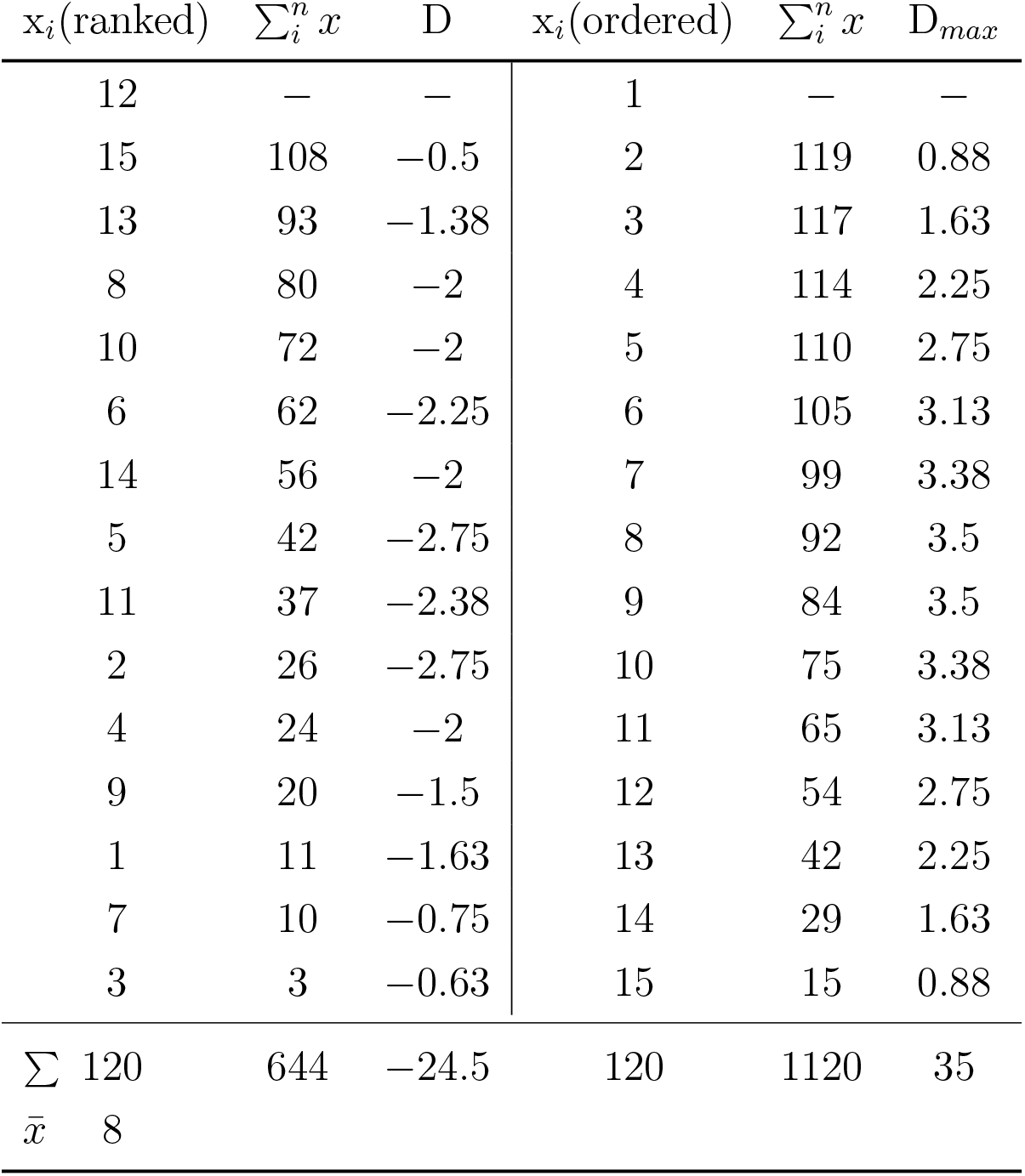
Evaluation of proportional differences.

## A new interpretation of Spearman’s rho

The abundance values in the example above have been chosen such they can alternatively can represent ranks. The reason for this is that the method used to assess univariate fluctuations of abundances over time can also be used to obtain the most widely accepted rank order correlation. Spearman’s Rho is give by:

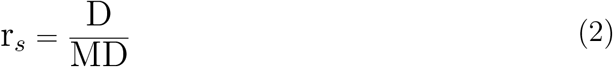

The ranks of one rank series are ordered from lowest to highest, D is then calculated for the corresponding values of the other rank series. It does not matter how the roles of the ordering and ordered rank series are assigned. However, there are slight differences when the approach is used to calculate the rank order correlation. If there are ties the familiar technique of taking the mean of the tied ranks is used in the ordered rank series. But ties can also exist in the ordering rank series, therefore the corresponding ranks in the ordered rank series are also averaged. For example, if the 3d and 4th ranks in the ordering series are tied and the corresponding values in the series that has been ordered are 2 and 8, then both of these values would have to be replaced by 5. The method yields the same values as the standard approach to correct the value of Spearman’s correlation which goes back to an attempt by Kendall(Taylor, 1994). The conventional method relies on calculating the sum of correction terms of the form 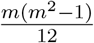 for each group of tied ranks with *m* tied values. This sum is then added to the sum of squared differences between ranks. The new approach is more straightforward and works also if more than 2 values are tied or if tied sets in both rank series overlap. Another difference to the calculation of V is also related to ties, MD used for standardizing D is not calculated from ordered ranks, which would include the tied ranks. Instead, the true maximum is calculated for the sequence of the first n natural numbers, i.e. by ignoring ties. Furthermore, it is not even necessary to calculate the maximum from proportional differences. The value of MD for n ranks can simply be calculated with this formula:

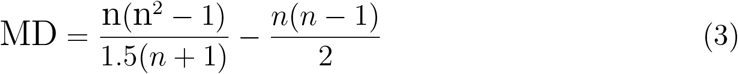

Overall, the new method makes the rank correlation easier to interpret, as it corresponds to the standardized sum of proportional differences between the mean rank and *n* – 1 subsets of ranks(2nd to highest, 3d to highest,…, highest) if one series is ordered according to the other series. The straightforward accommodation of tied ranks is another advantage over the conventional method.

## Evaluating trends in presence and absence

In certain ecological contexts and for certain taxa, population abundances are measured as binary data that only assumes one of two values, i.e. “presence” and “absence”, where absence is coded as 0 and presence as 1. A strong positive trend can reflect increasingly consistent use of a site and a strong negative trend that a site has been abandoned.

For the example in Table 2, trend measure V is given by 45/60 = 0.75. Without adjustment V equals 1 (or −1) as long as the observations are ordered. Unbalanced data (e.g., a far larger proportion of zeros than ones) is common in ecological datasets. To let V only indicate full correlation if the data is balanced it is possible to calculate the adjusted measure V_*b*_ by using a correction factor. The observations not only have to be in perfect order, the numbers of observed presences and absenced also has to be as equal as possible in order for V_*b*_ to assume the value of 1(or −1). If the number of time steps *n* is even, then V_*b*_ is given by:

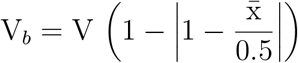

If *n* is an odd number, the following formula applies:

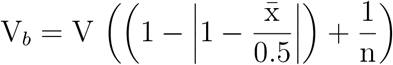

Here 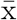 refers to the arithmetic mean of the observations that are coded as 0 and 1. For an even number of observations, the mean equals 0.5 if the data is balanced(i.e. equal number of ones and twos), and *V_b_* will have the same value as V. For a site with continuous absence for 14 time steps and then a single presence at the last time step V =1, but V_*b*_ = 0.20, as the correction factor in that case is 0.2. The same value with a negative sign is obtained for a time series with continuous presence for 14 time steps and absence at the last time step. For the example in Table 2 that has 8 observations without presence and 7 with absence the correction factor is 1 and V_*b*_ equals V. A high negative value would reflect a site that was used continuously in the past but has been abandoned just as long as it was used before. A high positive value that the site was not used in the past but has continuously been used in the second half of the observation period.

**Table 2.**
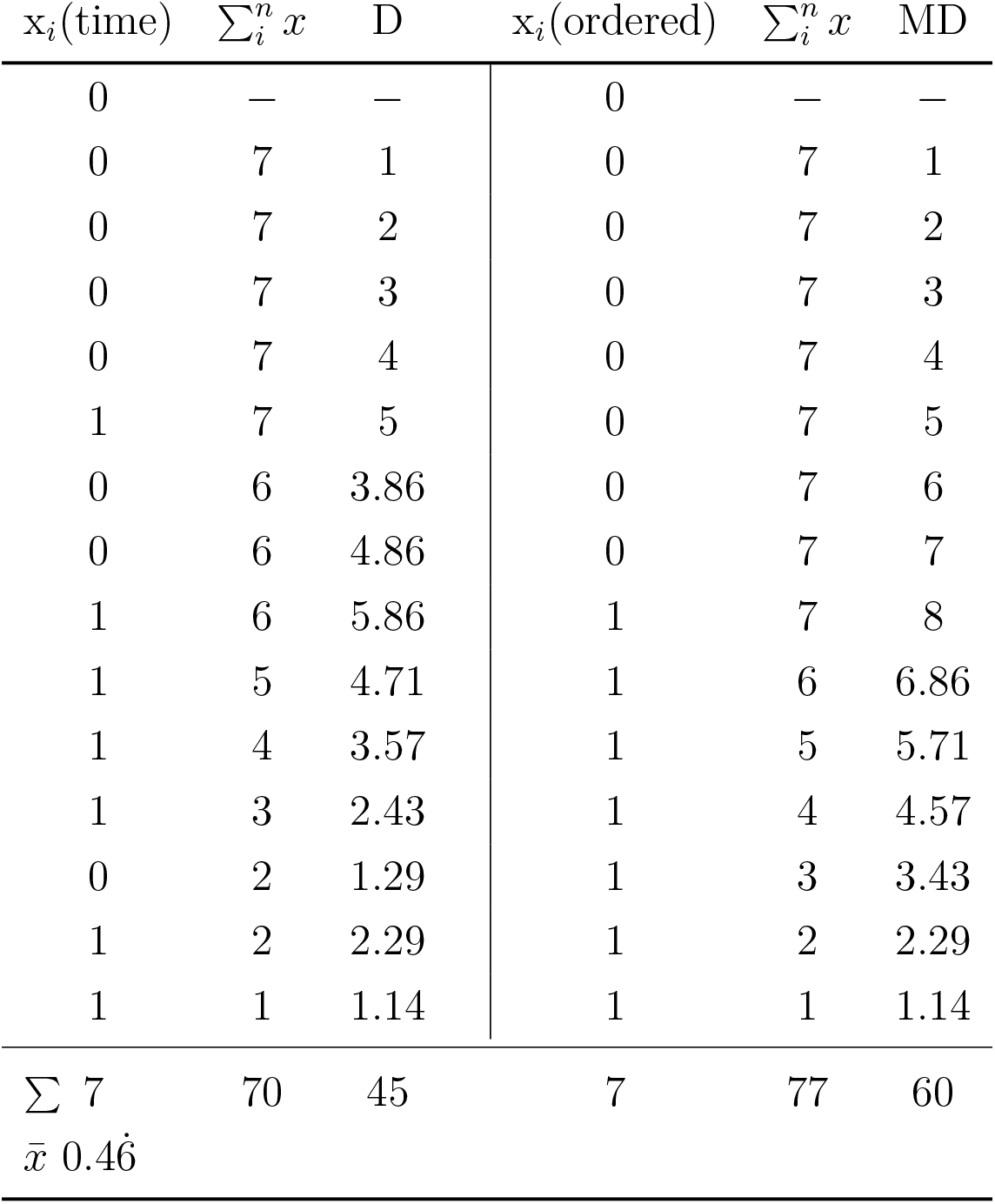
Evaluation of proportional differences.

## Sensitive rank correlation

The relationship between abundances of different species and between abundances and habitat variables is another active research area(Vázquez et al.,2007).To measure the bivariate relationship between variables correlation coefficients are used. The two most accepted measures of rank order correlations are Spearman’s Rho and Kendall’s Tau (Kendall, 1948, Lovie, 1995). Both these measures quantify the degree of the monotonic relationship between two rank series. A limitation of these rank correlations is that they lack sensitivity to top ranks (first, second, and so on). In many ecological contexts, top ranks can be considered to be more important than lower ranks, e.g. two conservation ecologists ranking the same list of individual species according to their importance for an ecosystem. Various weighted rank correlations have been described in the literature that extend existing measures to make them more sensitive to top ranks(see, Dancelli, 2013). Approaches of this kind are based on weights and there is often no justification for choosing a certain weighting scheme over another. A more straightforward approach would be to use a measure that is inherently more sensitive to top ranks and not an extension of an existing measure. Such a measure can be calculated from proportional differences between ranks.

**Figure 2:**
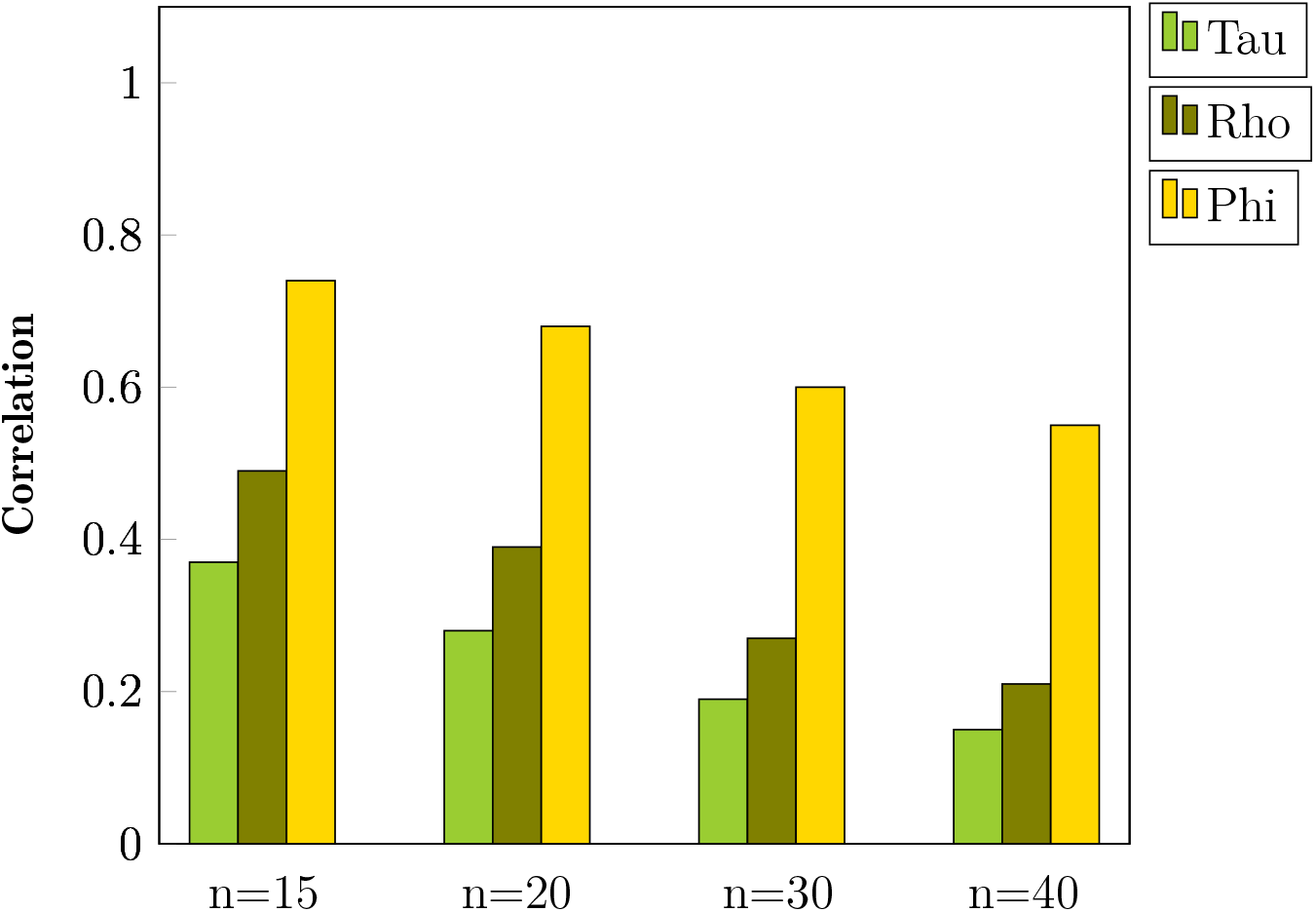
Comparison of the sensitivity to top ranks for Kendall’s Tau, Spearman’s Rho, and Phi. Each simulation was conducted 10.000 times for rank series of different lengths, with consistent concordance for the top 3 ranks, while the rest of the ranks for each bivariate distribution was generated randomly and without ties. Values correspond to mean correlations.

It is calculated as follows: First, one rank series is ordered according to the ranks of the other series, from lowest to highest. Then proportional differences relative to higher and lower ranks are computed for each rank. To calculate the difference to later time steps the sum of ranks that come later in the rank series is divided through the rank value, subtracting the number of comparisons yields the proportional difference, i.e. how often the value of the particular rank would need to be multiplied to fill the positive or negative difference to ranks that come later in the series. If the rank is on average lower than ranks that come later in the rank series this contributes a positive value to the proportional difference, denoted PD, if it is higher a negative value. Similarly, each rank is compared to the sum of ranks that come earlier in the rank series. Here the contribution to PD is positive if the rank is higher than ranks that come earlier, and negative if it is lower. To standardize PD the sum of all proportional differences is divided through the maximum, denoted MPD, that corresponds to the sum of proportional differences for a series that is ordered from lowest to highest, and that has no ties. Because the method yields different results depending on how the roles of ordering and ordered rank series are assigned, proportional differences are calculated twice, once for each rank series that is ordered by the other rank series. The resulting rank order correlation Phi can be written as follows:

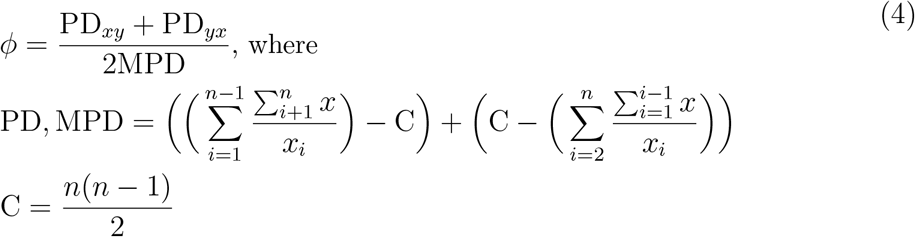

Phi is bounded between −1 and 1, reached only if all ranks are concordant or maximally discordant. The measure uses the whole range of values effectively. If there are tied ranks the approach described above in the context of Spearman’s rank-order correlation can be used. I conducted simulations for rank series of different lengths to assess the sensitivity of Phi and two standard measures of rank-order correlation toward top ranks. As shown in Figure 3, the new measure Phi indicates a higher correlation if there is an agreement between the top 3 ranks of both rank series compared to Spearman’s Rho and Kendall’s Tau. The influence of this concordance also decreases slower as *n* increases. Similarly, if there is discordance between top ranks of two rank series the measure would indicate a stronger negative association than standard rank correlations. But Phi can also indicate a lower correlation than the other measures, particularly if medium and lower ranks indicate a certain association that is contradicted by top ranks. These findings suggest that Phi is a rank order correlation that is inherently sensitive to top ranks.

**Figure 3:**
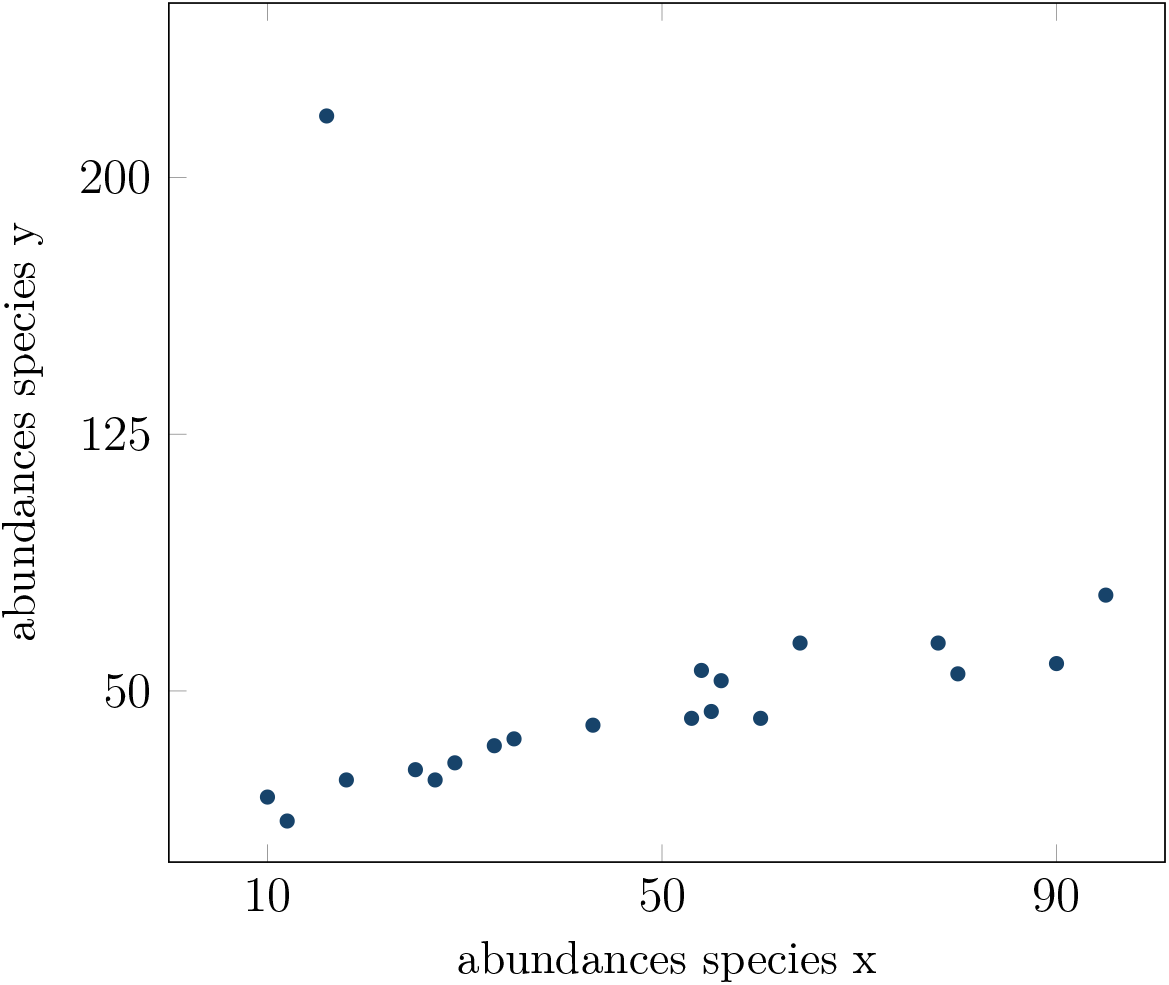
Example data with one outlier, r_*p*_ = 0.10 and Psi= 0.43

## Robust correlation for continuous variables

The degree of association between two variables is measured by a correlation coefficient. If both variables are metric the Bravais-Pearson product-moment correlation, denoted as r_*p*_, is used. Although the reduction of continuous data to ranks can lead to a loss of information, it is standard practice to use rank order correlations for originally continuous data if the assumptions underlying r_*p*_ are violated. These assumptions include bivariate normality, linearity, and the absence of outliers. The proportional approach can also yield a measure of bivariate association that does not have such limiting assumptions. The same differentials that underly trend measure V are used(see Equation 1), but each variable is ordered according to the abundances of the other variable. In this respect, the measure is similar to Phi(Equation 4). To differentiate the two proportional differences relative to the respective mean abundances they are denoted as D_*xy*_ and *D_yx_*, the corresponding maxima as MD_*xy*_ and MD_*yx*_.The measure of association, referred to as Psi, is then given by:

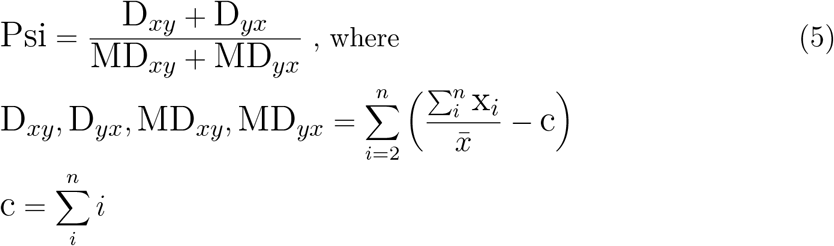

Psi takes on values in the range[−1, 1]. The measure can also be used if the data contains zeros or negative values, to avoid indetermination if the mean equals zero a small constant like 1 × 10^−9^ can be added to the mean. If there are ties in the variable that is used to order the other variable the same approach outlined above in the context of rank correlations can be used. I conducted simulations that revealed that Psi yields similar values to the Pearson’s correlation if values are drawn randomly from a normal or uniform distribution. But Psi has several advantages over r_*p*_. One assumption of r_*p*_ is that the relationship between variables is linear, that a change in one variable changes the other variable by a fixed amount. Conversely, the assumption underlying Psi is a monotonic relationship between variables, if one variable increases (or decreases), the other variable also increases (or decreases), and the value of one variable stays constant if the value of the other variable stays constant. This is often a more realistic assumption and a measure of monotonic association can provide a common ground for studies in variuos ecological contexts.

For r_*p*_ to be accurate, the data also cannot have outliers. A single extremely deviant observation can dramatically influence its value (Campbell, 2021). Such outliers typically reduce the absolute magnitude of the correlation coefficient and simply excluding them from the dataset is not considered legitimate(Schober, 2018). To cope with the problem of outliers and other violations of normality it has been recommended to use Spearman’s correlation, or conceptually similar methods that entail the transformation of the metric data to ranks, from which the Pearson correlation is then calculated(Bishara & Hittner, 2012). Figure 3 illustrates the effects of an outlier that illustrates the effect of a one-time population bonanza on measured correlations, Pearson’s correlation indicates a very weak positive correlation while Psi is more robust and indicates a moderate correlation. Psi is also more robust if there are low outliers, that correspond to population crashes. Therefore Psi is a viable alternative if there are outliers in the data, as long as they are not too extreme.

Attempts to apply certain cut-offs to describe the strength of the correlation can also be used for Psi and Phi. A classical resource on this approach is Evans (1996), and Evans’s standards for correlation strength are as follows: .00 to .19 is *very weak*, .20 to .39 is *weak*, .40 to .59 is *moderate*, .60 to .79 is *strong*, and .80 to 1.00 is *very strong*. Naturally, the rules apply invariably to positive and negative values of the new measures. The possibility of hypothesis testing with these measures remains to be investigated.

## Code availability

Excel workbooks and Jupyter Notebooks with code used for analyses are available at https://github.com/FT81/better-trend

## Notes

https://github.com/FT81/better-trend

### Competing Interest Statement

The authors have declared no competing interest.

### Summary of Updates

Figure added

